# Antiviral protection in the Pacific oyster *Crassostrea (Magallana) gigas* against OsHV-1 infection using UV-inactivated virus

**DOI:** 10.1101/2023.11.24.567680

**Authors:** Benjamin Morga, Mickäel Mège, Nicole Faury, Lionel Dégremont, Bruno Petton, Jean-François Pépin, Tristan Renault, Caroline Montagnani

**Author notes:** **Correspondence:** Caroline Montagnani, Benjamin Morga.

## Abstract

The increase of the frequency and severity of marine diseases affecting farmed marine mollusks are currently threatening the sustainability of this aquaculture sector, with few available prophylactic or therapeutic solutions. Recent advances have shown that the innate immune system of invertebrates can develop memory mechanisms allowing for efficient protection against pathogens. These properties have been called innate immune memory, immune priming or trained immunity. Previous results demonstrated the possibility to elicit antiviral immune priming to protect Pacific oysters against the ostreid herpes virus 1 (OsHV-1), currently plaguing *M. gigas* production worldwide. Here, we demonstrate that UV-inactivated OsHV-1 is also a potent elicitor of immune priming. Previous exposure to the inactivated virus was able to efficiently protect oysters against OsHV-1, significantly increasing oyster survival. We demonstrate that this exposure blocked viral replication and was able to induce antiviral gene expression potentially involved in controlling the infection. Finally, we show that this phenomenon can persist for at least 3 months, suggesting the induction of innate immune memory mechanisms. This study unravels new ways to train the Pacific oyster immune system that could represent an opportunity to develop new prophylactic strategies to improve health and to sustain the development of marine mollusk aquaculture.

## 1 Introduction

The Pacific oyster *Magallana gigas* (formely known as *Crassostrea gigas)* represents one of the main marine invertebrate aquaculture species in the world and in France, which plays a significant role in economy and in ecology of coastal areas. However, the increase of the frequency and severity of marine diseases currently threatens oyster production worldwide (1). Since 2008, Pacific oysters are plagued with high recurring mortalities worldwide sometimes identified under the term Pacific oyster mortality syndrome (POMS). This syndrome is a polymicrobial disease triggered by a variant of the ostreid herpes virus 1 (μVar) that is specifically affecting spats on oyster farms, although all stages, from larvae to adults, are susceptible to the virus when oysters are naïve (2-4). This syndrome has a significant impact on production due to the high mortality rate of OsHV-1 infected animals with up to 100% mortality within days (5). To date there is no existing prophylactic or therapeutic treatments available to mitigate mortality events (3). Existing solutions based on genetic selection strategies have been shown to improve Pacific oyster resistance to OsHV-1 infection. However, this strategy could impact oyster genetic diversity, with tradeoffs that might jeopardize their resilience potential against future diseases (6, 7). Solutions for biosecurity for viral pathogens has also been explored using seawater treatments that could be applicable in closed hatchery systems but still leave susceptible young oysters vulnerable when placed in open sea (8, 9). Current cultural practices also consist in the immersion in the environment of larger spat oyster quantities to cope with losses due to massive mortality events. A practice where extensive quantities of dead and sick spats are left to decay in open sea is not without consequences on the environment and could favor the spread of diseases (10-12). Altogether these control strategies have been insufficient to durably fight OsHV-1 infections. In addition, some of them may constitute aggravating factors and a major limitation to ensure sustainable aquaculture development that calls for innovative ways to mitigate these diseases.

Over the past decade, mounting evidence has demonstrated that invertebrates were capable to implementing antibody-independent innate immunological memory named “immune priming” (IP) or trained immunity. IP has been shown, in various invertebrate species that lack acquired immunity, to protect individuals against pathogens within-generation, across life stages (ontogenic IP) and even across generations (trans-generational IP, TGIP) (13, 14). Owing to their evolutionary significance and application perspective, these adaptive characteristics have emerged as an important new property of innate host defense mechanisms. In mollusks, there is an accumulating number of studies that evidenced forms of IP (see for review (15) (16)). In marine gastropods and bivalves, a few studies have investigated IP towards bacterial infections, in abalones, scallops, mussels and in *M. gigas* oysters. They collectively show that enhanced immunity can be induced following *Vibrio* bacteria pre-exposure (17-21). Regarding antiviral protection, to date, antiviral IP has been explored in a handful of studies on invertebrates (22, 23). Most studies were focused on crustaceans where several studies reported the potential of vaccine-like approaches using live, inactivated viruses (or derived elements) to fight major viral pathogen (24). In mollusks, we showed that a viral nucleic mimic (synthetic double stranded RNA called poly(I:C)) could induce a 100% protection for up to 5 months against OsHV-1 (25, 26). Exposure of oysters to poly(I:C) induced anti-viral IP in a dose-dependent manner and could be delivered either through injection or mucosal exposure (bath treatment). This protection was shown to be supported of a sustained antiviral state blocking viral replication in Pacific oysters, mediated by conserved candidate molecules and pathways (AMP, pattern recognition receptors, DSCAMs, FREPs, RNAi, Toll and IFN pathways) (27, 28). However, these previous works have limitations based on the very use of poly(I:C) such as economical cost, ethical and safety issues for human consumption. These studies nonetheless constitute a major breakthrough and open the way to design innovative strategies to overcome recurrent oyster diseases.

In this study, we investigated an IP-driven methods to protect oysters against OsHV-1 infection using UV-inactivated variant μVar. Numerous strategies have been used to evidence IP in invertebrate species notably using inactivated bacteria but also viruses. UV treatment was proven to inactivate the ostreid herpes virus 1 in previous studies (29, 30). The objectives of the present study were to test experimentally (i) the impact of UV treatment of OsHV-1 infectivity, (ii) the impact of UV-inactivated OsHV-1 injection on the antiviral immune response of the oyster, and (iii) the protection potential of this treatment upon secondary viral infection (antiviral immune priming). We show that UV-treatment efficiently impairs OsHV-1 infectivity. The injection of UV-inactivated OsHV-1 induces an antiviral state and an effective immune priming protecting the oyster against the infectious virus.

## 2 Methods

### 2.1 Pacific oysters

For the experiments described in §2.5.1 and 2.5.2, *Magallana gigas* spats were produced at the Ifremer facilities in La Tremblade in March 2018. Two batches were used, named SC18 and H12, and for each, 30 oysters were opened and sexed. Gametes were collected by stripping the gonad. For females, all gametes were combined and then sieved using 20-μm and 100-μm screens to remove small and large debris, respectively. Eggs were then divided by the number of males, from which the sperm was individually collected. Eggs were fertilized with the sperm and, 10 min after fertilization, all eggs were combined and then transferred into 30 L tanks. This method reduces sperm competition and thus maximizes the effective population size (31). The methodology used for the larval and spat cultures is described in two previous studies (32, 33). The progenies were maintained in our controlled facilities until priming experiments in 240-L raceways (at 14 to 20°C) with a continuous UV-treated and filtered seawater flow and an *ad libitum* phytoplankton diet (*Isochrysis galbana, Tetraselmis suecica and Skeletonema costatum*).

For the experiments described in supplementary files 2 and 4, spat oysters named 01/2020 and F15 were used. The 01/2020 oysters were progenies of wild oysters (n = 140 parents) that originated from a farming area in Marennes-Oléron (France). They were produced on August 2019 as previously described (34). These oysters were maintained free of pathogens under controlled bio-secured conditions until the onset of the experiment. Analyses of these oysters showed the absence of OsHV-1 DNA detection by qPCR and a low level of *Vibrio spp*., ∼1 CFU 100 mg−1 tissues (n = 3 pools of 5 ind.) (35). Similarly, the F15 oyster family (G2) were produced on March 19, 2020 using 25 siblings (G1), which were produced from two wild oysters (G0) from the Brest harbor (3).

Before the experiment, spat were acclimated in 120 L tanks with a continuous flow of UV-filtered seawater heated at 19°C. The seawater was enriched with the same phytoplankton mixture. The acclimation process spanned at least 2 weeks to ensure optimal growth and feeding conditions, essential for observing effective virus replication leading to oyster mortalities (2).

### 2.2 Poly(I:C) preparation

Poly(I:C) HMW (Invivogen, cat. code:tlrl-pic) was injected following the same protocol as in Lafont et al., 2017 (26). Briefly, poly(I:C) was resuspended in sterile water to a final concentration of 1 mg/ml, 100 μL were injected, which resulted in each oyster been injected with 100 μg of poly(I:C).

### 2.3 Viral suspension preparation

Viral suspension was obtained following previously established protocol (29) from OsHV-1-tissue homogenate from contaminated oysters. Virus-free suspension was prepared following the same protocol but using OsHV-1-free oysters.

### 2.4 Inactivation of OsHV-1 by UVC

Ultraviolet (UV) light treatment for viral suspension and virus-free suspension was performed using low pressure UVs, (1 min and 45 seconds illumination under wave lengths of 254 nanometers corresponding to 0.354mW / cm2) monitored with a VLX-3W radiometer.

### 2.5 Experimental designs

#### 2.5.1 UV-treated OsHV-1 protection experiments

To investigate the impact of UV-treatment on OsHV-1 infectivity and the impact of exposition of UV-treated OsHV-1 on oyster susceptibility, the following experiment was performed (Sup. Figure1A): In a primary exposure, oysters (n=140/per condition, H12 population, 10 month old) were first injected either with (a) UV-treated OsHV-1 suspension (virus UV), (b) UV-treated virus free suspension as a negative control (free virus UV), (c) Poly (I:C) as a positive control (PIC) and (d) filtered seawater as a negative control (FSW). 24hrs after primary exposure, each treatment group (a to d) was separated in two groups of seventy oysters that were exposed either to contaminated seawater with OsHV-1 (CSW) or non-contaminated seawater (SW), *i*.*e*. 8 conditions in total: virus UV/CSW; virus UV/SW; no virus UV/CSW, no virus UV/SW; PIC/CSW; PIC/SW, FSW/CSW; FSW/CSW.

The protocol of infection by OsHV-1 μVar was based on immersion into titrated contaminated seawater. Briefly, 100 healthy oysters were injected with 100μL of OsHV-1 suspension (1,37.10^7^UG/μL) and placed in a 30L tank (temperature 20°C). After 24 hours, the oysters were removed from the tank and the contaminated seawater was dispatched to different aquaria. Forty oysters per condition were bathed in 3L of contaminated seawater separated in two aquaria for sampling. Thirty oysters were placed in two others aquaria for mortality monitoring. Mortality was recorded every day for 7 days in all conditions. Oysters were sampled (whole animals snap-frozen in liquid nitrogen, 10 oysters per condition per time point) before the primary exposure (T0), 24hrs post-primary exposure (T1), 24hrs post-secondary exposure (T2) and 48hrs post-secondary exposure (T3). This experiment was repeated at least once (Sup.Figure 1B).

**Figure 1.**
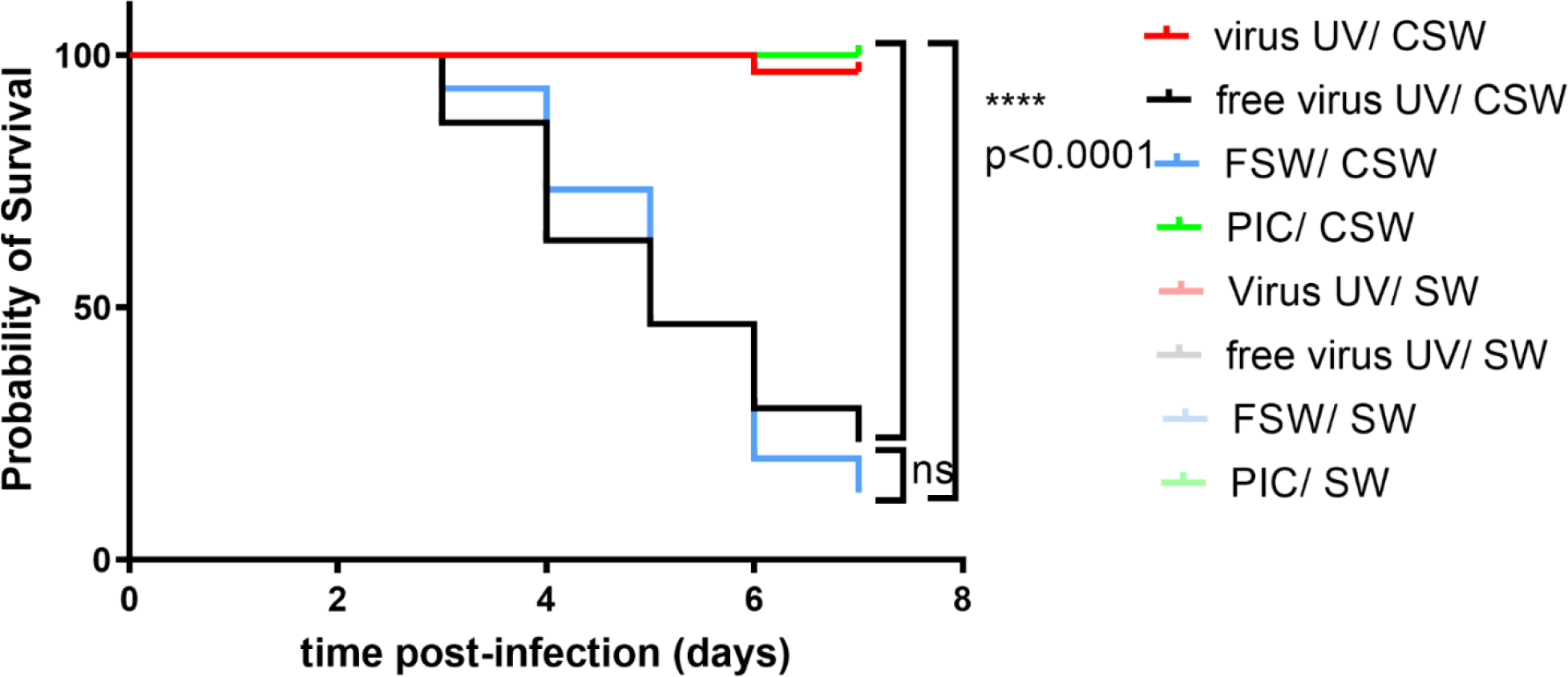
UV-inactivated OsHV-1 treatment increases oyster survival after virus infection. Kaplan-Meier curves showing probability of oyster (H12) survival after a primary exposure to UV treated OsHV-1 (virus-UV in red), poly (I:C) (PIC, in green), UV-treated virus free suspension (free virus UV, in black) or FSW (blue) and a secondary exposure to OsHV-1 contaminated seawater (CSW) (plain lines) or filtered seawater (FSW, faint color lines). Controls reaching 100% survival (non-treated control lines and oysters challenged with FSW appear hidden and merged behind the poly(I:C) positive control). Mortalities in each group of 30 oysters (15 per tank) were monitored for 7 days after infection. **** indicates p-value <0.0001; ns indicates non significative diffrences, log-rank test (n=30).

#### 2.5.2 Duration of protection

To test the duration of the protection, we used a susceptible oyster family (SC18, 8 months old) (Sup.Figure 3A). In a primary exposure, oysters (n=45/per condition) were first injected either with (a) UV-treated OsHV-1 suspension (virus UV), or (b) UV-treated virus free suspension (free virus UV), and (c) filtered seawater as negative controls (FSW). At 8 days, 1 month, 2 months, 3 months after primary exposure (each treatment group (a to c) was separated in two groups that were challenged by injection of 100 μL (3,9.10^4^ UG/μL) of OsHV-1 suspension (“virus” condition, n=30) or injected with 100 μL of filtered artificial seawater as a negative control (“SW”, n=15). Mortalities were monitored for 7 days after challenge. This experiment was duplicated using the same protocol and oysters (sup. Figure 3B).

### 2.6 DNA extraction and real time PCR analysis for OsHV-1 DNA quantification

For each oyster, a piece of mantle and gills was sampled and DNA extraction was performed using QiAmp tissue mini kit® (QIAgen) according to the manufacturer’s protocol. DNA was diluted to a final concentration of 5 ng/μl. The detection and quantification of OsHV-1 DNA was carried out by real time PCR (36). Assays included a standard curve and a negative control.

### 2.7 RNA extraction and real time PCR analysis for immune gene expression quantification

Frozen oysters were homogenized by bead-beading (Retsch, Mixer Mill MM400) with a stainless-steel ball bearing and housing that had been pre-chilled with liquid nitrogen. Frozen oyster powder (20 mg) was homogenized in 1 ml TRIzol by vortexing 1 h at 4 °C. Prior to extraction, insoluble materials were removed by centrifugation at 12,000× g for 10 min at 4 °C. Total RNA was extracted using TRIZOL® Reagent™ (Ambion®) according to the manufacturer’s recommendations. Total RNA was treated with Turbo™ DNAse (Ambion®) to remove genomic DNA. A second RNA extraction using TRIZOL was carried out and completed by RNA purification with Direct-zol RNA Miniprep (Zymo Research). RNA quality and quantity were determined using a NanoDrop 2000 (Thermo Scientific). First-strand cDNA synthesis was performed using the SuperScript® III First-Strand Synthesis System (Invitrogen) with 500 ng of RNA used. A No RT (No Reverse Transcription) control was performed in order to control absence of oyster genomic DNA using oyster elongation factor alpha (EF1 alpha) primers. For qPCR analyses, pipetting and amplification were performed with a Labcyte Acoustic Automated Liquid Handling Platform (ECHO) and a Roche LightCycler 480, on the qPHD -Montpellier genomix platform. The total qPCR reaction volume was 1.5 μl and consisted of 0.5 μl of cDNA (dilution 1/12) and 1 μl of SensiFAST™ SYBR® No-ROX Kit (Bioline) containing 0.4 μM of PCR primer (Eurogenetec SA)(primer list in sup. table 1). The amplification efficiency of each primer pair was validated by serial dilutions of a pool of all cDNAs.

Gene relative expression was calculated in two steps: First, the comparison ratio of a target cDNA sequence to a reference DNA sequence (mean of two housekeeping genes, *Cg*-RPL40 for ribosomal protein L40, Genbank AN FP004478; *Cg*-EF1 for elongation factor 1α, Genbank AN AB122066) was realized automatically by LightCycler® 480 Software (Version 1.5). Then, normalized ratio was calculated by dividing the ratio of selected condition with the ratio of control condition according to Pfaffl, 2001 (37):

Normalized ratio = ([target]/[reference]) selected condition/([target]/[reference]) control condition. For every condition, each of the 10 individuals at time T0 was compared to the 10 individuals at time T1 that generated 100 values/condition/gene.

### 2.8 Statistical analysis

Data were analyzed by using GraphPad Prism version 8 software (La Jolla California USA). Survival rates were analyzed by log rank test on Kaplan-Meier survival curves. A multiple t-test was performed to compare relative expression for each target gene between one time point and the control time. These statistical tests can be used without considering the data normality due to the high numbers of data (n=100). A Kruskal-Wallis complete by a post hoc Dunn’s Multiple Comparison test were performed to compare relative expression for each target gene from one control condition (FSW) in comparison to the three others conditions between time T0 (before priming, *i*.*e*. naïve oyster) and time T1 (24h post-priming).

## 3 Results

### 3.1 UV-inactivated OsHV-1 injection induces an antiviral protection in the Pacific oyster

We tested the impact of inactivated OsHV-1 exposure on oyster survival against the virus by injecting either UV-treated OsHV-1, a UV-treated oyster tissue homogenate (virus free) or filtered seawater (FSW) as negative controls, or poly(I:C) as a positive control (Figure 1), 1 day before infecting the oysters with infectious virus. Challenge was performed using natural routes of infection using contaminated seawater (CSW). Survival curves after challenge show that all conditions had a 100% survival after secondary exposure to filtered seawater, the negative control of infection. Conversely, oysters that were primary exposed to FSW (blue) or “free virus UV” controls (black) exposed to CSW reached low survival rates with 13% and 23% survival 7 days post-infection, respectively. However, oysters first exposed to UV-treated OsHV-1 (10^4^ copies/μL) demonstrated significant (p<0.0001) higher survival rates than the control (FSW-CSW) with 97% survival 7 days post-infection. The optimal dose of UV-treated OsHV-1 was established in parallel experiments (sup. Figures 2A,Ba,b). These results were similar to the priming control where oysters were firstly exposed to poly(I:C) and subsequently infected with CSW (no significant differences, 100% survival). These results were also confirmed in a replicate experiment (sup.Figure 1B).

**Figure 2.**
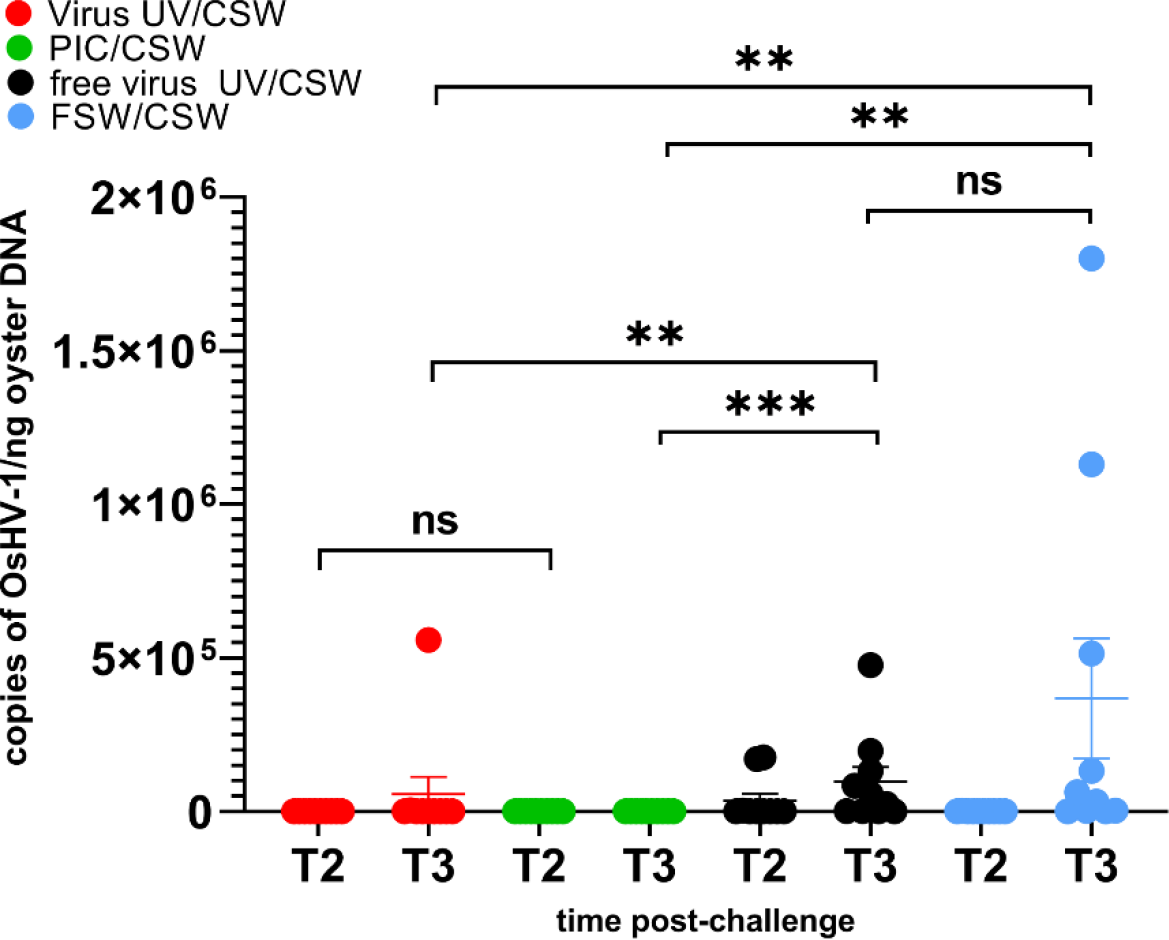
UV-inactivated OsHV-1 treatment blocks viral replication. Viral DNA loads after challenge with live OsHV-1 were estimated by real time PCR in 10 individual oysters for each condition (UV inactivated virus in red, UV-treated free virus suspension in black, filtered seawater (FSW) in blue, poly(I:C) in green) 24h (T2) and 48h (T3) after challenge with contaminated seawater (CSW). DNA loads are expressed as copies of OsHV-1/ ng of oyster DNA. Mann-Whitney T test, ** pvalue < 0,05; *** pvalue<0,001, ns-non significative (n=10).

**Figure 3:**
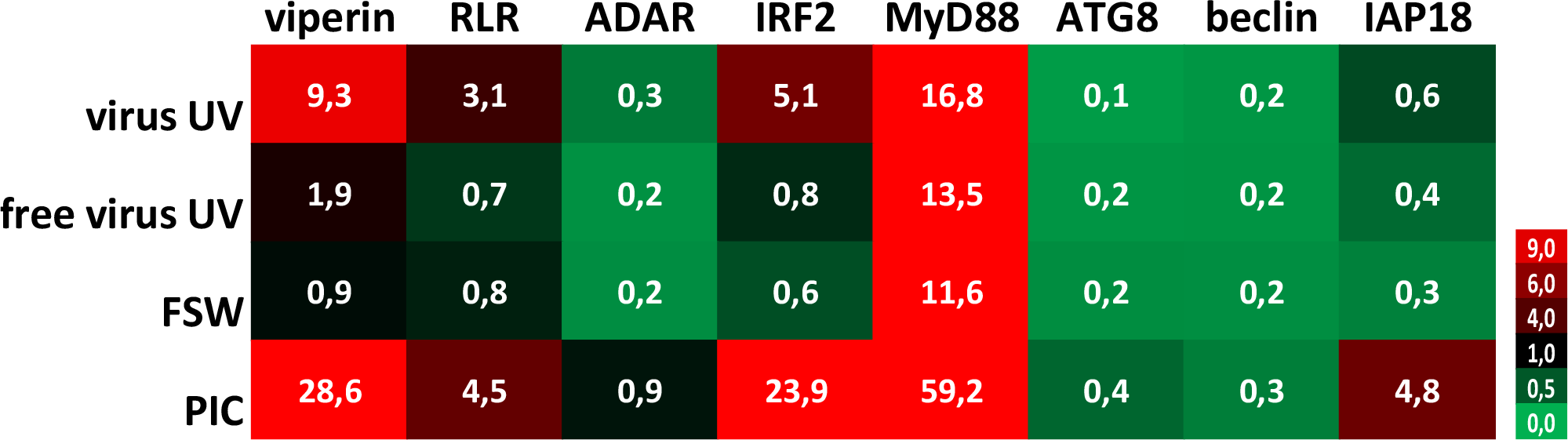
UV-inactivated OsHV-1 treatment induces antiviral gene expression. Heatmap of the expression level of 8 candidate genes. Each value corresponds to comparisons of time T0 (naïve oysters) with time T1 (T24hrs post-primary exposure) mean fold changes for each condition (n=100 values) (UV-treated virus = virus; UV-treated non viral suspension = free virus UV; filtered seawater =FSW; poly(I:C) injection = PIC).

The virus DNA amount in all conditions was quantified by following OsHV-1 DNA in the oysters 24h (T2) and 48h (T3) after infection (Figure 2). High virus DNA loads were observed in “free virus UV” and “FSW” conditions, reaching a mean of 9,7.10^4^ and 3,7.10^5^ genome unit/ng of DNA at 48hrs post-infection, respectively. We observed an evolution of the viral DNA load between 24 to 48 hours reflecting a viral replication in “free virus UV” and “FSW” conditions that was confirmed in virus gene expression analyses on three viral genes (sup.Figure 1C). These two conditions did not show any significant differences in viral DNA amounts at T3. Significantly lower DNA virus amounts (p-value <0.05) were detected in “PIC” and “virus-UV” conditions. Viral DNA loads in poly(I:C) and UV-treated virus conditions were not significantly different 24hrs after infection. More elevated viral loads were observed in the UV-treated virus and poly(I:C) at T3 with a mean of 1,73.10^2^ and 5,6.10^4^ genome units/ng DNA, respectively. This observation might result from the sampling of animals non-responsive to the treatment (corresponding to the 3.3% mortality rate at the end of the experiment) and/or suggests that the treatment is still progressing in blocking viral replication. In accordance, complementary results where viral DNA loads were quantified in surviving oysters 7 days after challenge revealed low or undetectable virus DNA in oysters exposed to this dose of UV-treated OsHV-1 (sup. Figure 2Bc, d).

### 3.2 UV-inactivated OsHV-1 injection induces antiviral gene expression stimulation

To evaluate the impact of UV-treated OsHV-1 injection on the oyster antiviral response, we monitored the expression of 8 candidate genes that were chosen for their implication in antiviral response in the oyster as decribed in previous studies (sup.table 1). These genes are belonging to the Interferon-like and NF-κB pathways (from receptor-RLR-, to signal transducers -MyD88-, transcription factors-IRF- and antiviral effectors-ADAR, viperin) or apoptosis -IAP-, or the autophagy pathway -ATG8, Beclin) (27, 38-40). To illustrate changes in antiviral immune gene expression according to treatment (primary exposures to UV-treated virus, no virus UV, FSW and PIC), a heatmap (Figure 3) was generated to visualize the variability of the expression level of the 8 candidate genes. This heatmap represents comparisons of mean fold changes for each condition between time T0 (naïve oysters considered as non infected) and time T1 (24hrs post-primary exposure). It clearly shows that UV-inactivated virus injection induced an upregulation of antiviral genes except for ADAR and autophagy-related genes. For all gene tested, except for viperin, no significant differences were observed between the “free virus UV” and “FSW” conditions (sup.Figure 1D). Significant differences were observed between these control conditions and either “UV-virus” and “PIC” conditions (sup.Figure 1D). However, this induction was lower in the “UV-virus” condition than in poly (I: C). Poly(I:C) was the most efficient in inducing gene up-regulation, even for the IAP gene that didn’t seem to be regulated in the other conditions.

### 3.3 UV-treated OsHV-1 treatment induces long-term protection in M. gigas against OsHV-1 infection

To test whether this observed protection could last over 24hrs, we expanded the lapse time between UV-treated OsHV-1 injection and OsHV-1 challenge. To this end, oysters were exposed either with UV-treated OsHV-1, UV-treated virus free tissue homogenate or with FSW as controls and stored in a bio-secured nursery facility (sup.Figure 3A). Oysters from the different conditions were then challenged by injection with an OsHV-1 suspension prepared from the same batch of infected oysters, 8 days, 1 month, 2 and 3 months after treatment. Kaplan-Meier survival curves at all time points showed that UV-treated OsHV-1 was still efficient in protecting oysters from 24h to 3 months post-priming with significantly higher survival rates (p<0,0001) ranging from 100% survival at day 8 to 93%, 87% and 63% after 1, 2 and 3 months, respectively (Figure 4), corresponding to a gain of survival due to the priming of 100, 83, 77, and 47% respectively. Those results were confirmed over two months in a replicate of this experiment (sup.Figure 3B).

**Figure 4.**
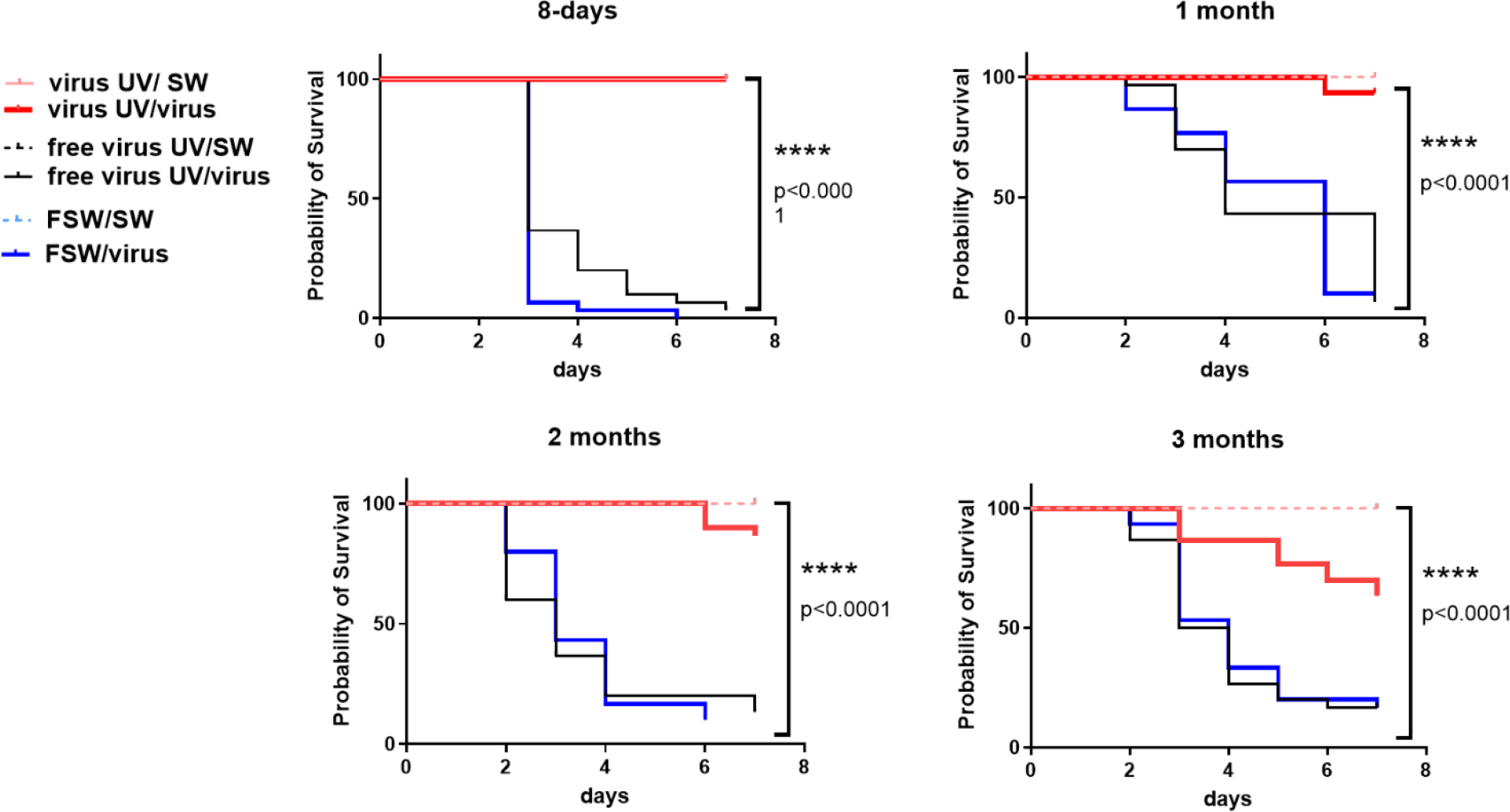
UV-inactivated OsHV-1 treatment induces a long-term protection against OsHV-1 infection. Kaplan-Meier curves showing probability of oyster survival after a primary exposure to UV-treated OsHV-1 (virus-UV in red), UV-treated non-viral suspension (free virus UV, in black) or filtered seawater (FSW in blue) and a secondary exposure to OsHV-1 contaminated seawater (CSW) (plain lines) or filtered seawater (FSW, dotted lines), 8 days, 1 month, 2 months and 3 months after the primary exposure. Controls reaching 100% survival (oysters challenged with FSW) appear hidden and merged behind the non-treated control line. Mortalities in each group of 30 oysters (15 per tank) were monitored for 7 days after infection. **** indicates p-value <0.0001; log-rank test (n=30).

## 4 Discussion

In this study, we investigated the use of UV-inactivated OsHV-1 to induce immune priming as an alternative method to poly(I:C) to protect Pacific oysters from viral infection. UVC (200-280nm) irradiation is known to inactivate viruses by chemically modifying their genetic material but leaving the overall structure of viral protein complexes mostly intact (41, 42). Although other treatments have been shown to inactivate viruses, we chose this treatment because it is possible to obtain viral material that retains immunogenic properties of the native antigens. Our results show an improved survival rate in animals primary injected with UV-treated viral suspension (from 96.7% to 100%) and with poly(I:C) (100%) as opposed to animals primo-injected with seawater. The difference between the survival rates for the UV-inactivated virus injected animals and negative controls were highly significant in all experiments (log-rank test p-value <0.0001). Additionally, this protection was observed in at least three different genetic backgrounds suggesting that this phenotype is not dependent on a specific genetic immune capacity related to a specific Pacific oyster family.

Treatments using heat inactivation of OsHV-1 have recently been tested in oysters but demonstrating lower efficiency in protection against OsHV-1 infection (43). Moreover, this treatment was efficient in protecting the oysters over three months. Further experiments should investigate whether this protection could be enhanced, for example by repeating the exposition, and last for longer time periods as for poly(I:C) which was found effective for at least 5 months (26). The 3-month protection however suggests that the exposition to UV-inactivated OsHV-1 have induced immune priming mechanisms. We also showed that oysters primed with UV-inactivated virus, as for poly(I:C), have only traces of viral DNA or RNA, suggesting that the treatment and the induced immune status inhibit viral replication. This result also demonstrate that the virus inactivation was efficient and complementary analyses revealed that this inactivation was stable over time and at different temperatures with no evidence for reactivation of the virus that was undetectable in primed animals 7 days after challenge (sup.Figure 4).

To better understand this phenomenon, we investigated the impact of the priming on the expression of some oyster immune genes. We show that the UV-inactivated OsHV-1 is still able to induce and overexpression of immune r-related genes. This was especially true for the viperin, RLR, IRF2 and MyD88 genes which is consistent with previous studies where these genes were also implicated in the priming response to poly(I:C) (27). This up-regulation was significantly different from the one observed in oysters injected with filtered seawater or a UV-treated oyster tissue homogenate free of virus. Viral gene induction was also significantly lower in the UV-inactivated condition that in the poly(I:C) which might suggest distinct responses that could be triggered by distinct viral features.

Altogether, these results hold promises to develop IP-driven methods to mitigate with recurrent bivalve mollusk diseases. However, further investigations will be required to develop this method and evaluate potential risks associated with it. Several issues will have to be addressed as the optimization of the method to adapt to large-scale application in Pacific oyster farms. Future applications in aquaculture would greatly benefit from an early treatment when batch exposition is more practical, such as in larvae or post-metamorphosis stages. Additionally, transgenerational immune priming (TGIP) represents an opportunity for implementation of IP strategies in cultivated and hatchery-produced Pacific oysters. TGIP has been so far reported in a dozen invertebrate species, sometimes for several generations (13). Recent data also suggests that poly(I:C) or commensal microbiota exposure can induce immunological memory that persists across generations (44, 45). Priming the genitors could be an efficient way of producing millions or billions of progenies that can display enhanced immune capacities.

Another issue would be the impact of the treatment on oyster fitness. Caution should indeed be taken, as we currently do not know if trade-offs could occur between enhanced immunity and other important physiological or desirable production traits in oysters. Persistent immune protection and an overall increased capacity of host defense could be beneficial in long-lived organisms, likely to face repeated exposure to the same or similar pathogens. However, this functional investment may have damaging effects if activated at the wrong time and in an inflated fashion. In vertebrates, it has been suggested that IP may play a role in the pathogenesis of auto-inflammatory and/or autoimmune diseases (46). In invertebrates, previous studies have already emphasized the potential trade-offs linked to IP notably on nutrient-demanding processes such as reproduction, growth, and possibly other life-history traits (47). A previous study already revealed potential costs of poly(I:C) priming, notably on larval development, that could have deep implications on oyster fitness and would then require further investigations (44, 48). In this study, this induction of immune genes by the inactivated virus was lower than the upregulation observed when the oyster was facing poly(I:C). The fact this treatment induces an efficient protection with milder induction of antiviral genes than poly(I:C) could be advantageous for future applications in reducing the impact of the treatment on oyster physiology and limiting potential trade-offs on the long term.

Finally, the mechanisms behind this protection should be further investigated to fully understand the impact of the treatment on the immune system and the mechanisms supporting IP and immunological memory. Several studies have now suggested the participation of invertebrate immune cells in IP, notably in oysters (49). However, how IP information is recorded and stored is still an open question. Epigenetic reprogramming and the rewiring of intracellular metabolic pathways have been considered as one of the main mechanisms driving trained immunity and TGIP in mammalian innate immune cells, but also in several invertebrate species, or in plants (16, 50-53). Previous results already point towards a role of epigenome remodelling in immune memory formation and IP in *M. gigas*. Transcriptomic modifications following poly(I:C) IP indeed evidenced differential regulation of genes related to the epigenetic machinery (27). DNA methylation has also been identified as a support for immune memory in response to early larval microbiota interaction (45). The rapidly growing field of epigenetic and metabolomics studies should allow us to further investigate epigenetic support IP in oysters and whether it can target specific immune pathways and results from metabolic reprogramming of immune cells.

In conclusion, these results offer us today a unique opportunity to consider IP-driven methods to protect Pacific oysters against infection due to OsHV-1 and other pathogens resulting in a better control of related mortality events. This innovative solution (results protected under patent (54)) will help to develop prophylactic strategies and finally prevent the massive mortality events affecting this economically and ecologically valuable shellfish species and contribute to the sustainable management of this resource.

## 6 Author Contributions

CM, BM, NF designed experiments. BM, CM, MM, NF and LD performed oyster experiments. LD and BP performed oyster reproduction. TR and JFP performed inactivated OsHV-1 by UVC. BM, CM, NF and MM performed OsHV-1 DNA quantification and viral genes expression analyses. BM, CM, NF, MM interpreted results. CM, BM wrote the manuscript. All the authors have revised and approved the manuscript submission.

## 8 Acknowledgments

We acknowledge the support of the Ifremer facilities of La Tremblade, Bouin and Argenton (ASIM-“Adaption et Santé des Invertébrés Marins”, “Atlantic Marine Molluscs Experimental” -EMMA and PHYTNESS units, France). Authors also thank the FINA Ifremer program and the DECICOMP project (Guillaume Mitta, ANR-19-CE20-0004) for 01/2020 and F15 oyster production, respectively. We are grateful to Philippe Clair from the qPHD-Montpellier GenomiX platform for useful advice.

