## supplementary figures for "Antiviral protection in the Pacific oyster *Crassostrea (Magallana) gigas* against OsHV-1 infection using UV-inactivated virus"

### 1 Supplementary tables

#### 1.1 sup. Table 1. list of primers used for oyster gene and virus gene expression analysis

| Name | Organism | GenbankAN | sequence | | Function | Efficiency | Reference |
|---|---|---|---|---|---|---|---|
| *Cg*-EF1 | oyster | AB122066 | forward | GAGCGTGAACGTGGTATCAC | housekeeping | 2,1 | de Lorgeril et al., 2018 (3) |
| | | | reverse | ACAGCACAGTCAGCCTGTGA | | | |
| *Cg*-RPL40 | oyster | FP004478 | forward | AATCTTGCACCGTCATGCAG | housekeeping | 2,0 | de Lorgeril et al., 2018 (3) |
| | | | reverse | AATCAATCTCTGCTGATCTG | | | |
| viperin | oyster | EKC28205 | forward | TAAATGCGGCTTCTGTTTCC | IFN pathway | 1,9 | Lafont et al., 2020 (24) |
| | | | reverse | CAGCTGAAGGTCCTCTTTGC | | | |
| RLR | oyster | EKC34573 | forward | CCGGAAGTATGCTAGAGGAAGA G | IFN pathway | 1,9 | Lafont et al., 2020 (24) |
| | | | reverse | TCCGGTGTAGTTGTAGGTGATG | | | |
| ADAR1 | oyster | EKC20855 | forward | CTCAAACAGTGCAACTGCATC | IFN pathway | 2,0 | Lafont et al., 2020 (24) |
| | | | reverse | TCACAAGCCCTGCTATCAC | | | |
| IRF2 | oyster | EKC43155 | forward | CGAAACGCAGAAACTGTTC | IFN pathway | 2,0 | Lafont et al., 2020 (24) |
| | | | reverse | ATTTGCCTTCCATCTTTTGG | | | |
| IFI44 | oyster | XM_020065029 | forward | CACCAGTTTGGAAACCGCAG | IFN pathway | 2,0 | Lafont et al., 2020 (24) |
| | | | reverse | GATACTGGATTTTCCCGCGC | | | |
| MyD88 | oyster | DQ530619.1 | forward | CGTGCCATGGACGGATAACAACG | NF-kB pathway | 1,8 | Lafont et al., 2020 (24) |
| | | | reverse | GGCCCAGCAGTACCTCTGTGGAATC | | | |
| ATG8 | oyster | EKC40439.1 | forward | CCGATGCTTGACAAGACCAA G | autophagy | 2,0 | Picot et al., 2020 (36) |
| | | | reverse | CCGTCCTCGTCTTTCTCCTG | | | |
| Beclin | oyster | EKC28450.1 | forward | AAATGCTGCTTGGGGTCAG | autophagy | 2,0 | Picot et al., 2020 (36) |
| | | | reverse | CGGAATCCACCAGACCCATA | | | |
| IAP18 | oyster | JH818926 | forward | CCCGAAAACGTAACCTCAGA | apoptosis | 2,0 | Segarra et al., 2014 (37) |
| | | | reverse | TTTCGTTTGCTGCTCATTTG | | | |
| ORF80 | OsHV-1µVar | YP_024619.1 | forward | AAGAGGATTTGGGTGCACAG | Membrane protein | 2,0 | Morga et al., 2017 (38) |
| | | | reverse | TTGCATCCCAGGATTATCAG | | | |
| ORF87 | OsHV-1µVar | YP_024626.1 | forward | CACAGACGACATTTCCCCAAA | Apoptosis inhibitor | 2,0 | Morga et al., 2017 (38) |
| | | | reverse | AAAGCTCGTTCCCACATTGGT | | | |
| ORF99 | OsHV-1µVar | YP_024638.1 | forward | GGTGGAGGTGGCTGTTGAAA | Apoptosis inhibitor | 2,0 | Morga et al., 2017 (38) |
| | | | reverse | CCGACTGACAACCCATGGAC | | | |

### 2 Supplementary Figures

#### 2.1 Supplementary Figure 1

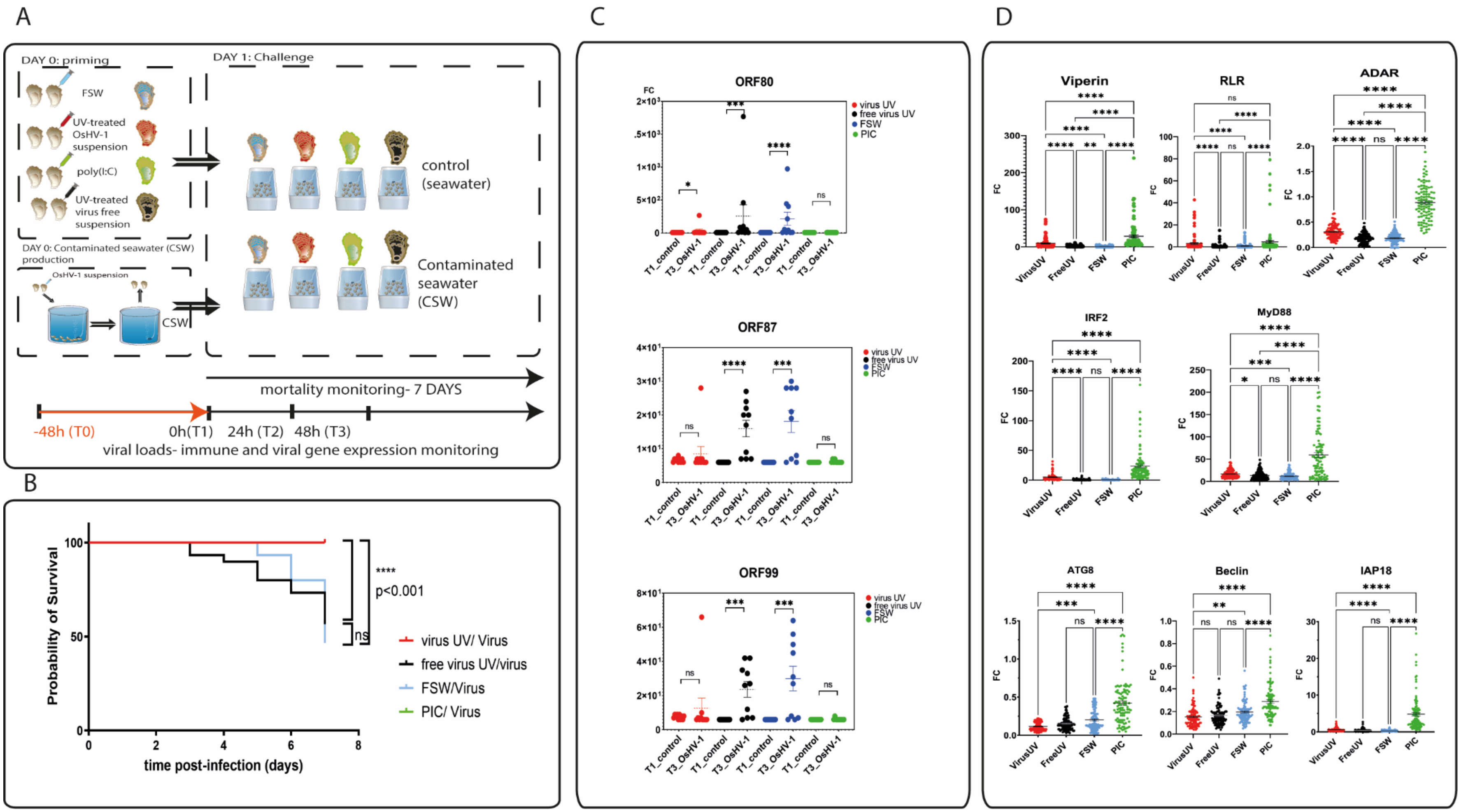


***Sup.Figure 1. Impact of treatment with UV-inactivated OsHV-1 on oyster survival and viral infection***

*A. **Experimental design for immune priming using UV-inactivated OsHV-1 suspension (mortality monitoring in Figure 1).** OsHV-1 susceptible oysters were first exposed by injection to UV treated OsHV-1 (virus-UV in red), poly (I:C) (green), UV-treated virus* *free suspension (in black) or filtered seawater (FSW in blue) one day before being challenged with contaminated seawater (CSW) or filtered seawater as a control. Mortalities were monitored daily for 7 days after the challenge and oysters sampled for viral DNA, RNA and oyster RNA extractions at T0 (beginning of the experiment), T1 (24hrs after priming), T2 (24hrs after challenge) and T3 (48hrs after challenge).*

*B.* ***UV-inactivated OsHV-1 treatment increases oyster survival after virus infection.*** *Kaplan-Meier curves of an independent experiment (replicate to Figure 1) showing probability of oyster survival after a primary exposure to UV treated OsHV-1 (virus-UV in red), poly (I:C) (PIC, in green), UV-treated non-viral suspension (free virus UV, in black) or filtered seawater (FSW in blue) and a secondary exposure to OsHV-1 contaminated seawater (CSW) (plain lines). Controls reaching 100% survival (poly(I:C)/virus) appear hidden and merged behind the virus-UV/virus line. Mortalities in each group of 30 oysters (15 per tank) were monitored for 7 days after infection. **** indicates p-value <0.0001; ns indicates non significative differences, log-rank test (n=30).*

*C.* ***UV-inactivated OsHV-1 treatment blocks viral replication.*** *OsHV-1 ORF80-87-99 expression was monitored using the same protocol as for oyster gene monitoring in samples from experiment described in sup.Figure1. Gene expression was compared in oysters sampled 24hrs after priming (T1, before challenge) and 48hrs after challenge (T3) with CSW for the four treatment conditions ("virus UV" in red, "free virus UV" in black, FSW in blue and poly(I:C) in green). Mann-Whitney T test, * pvalue < 0,05; *** pvalue<0,005, ****pvalue<0,0001, ns- non significative (n=10).*

*D.* ***UV-inactivated OsHV-1 treatment induces antiviral gene expression.*** *Oyster gene expression was monitored using qPCR in samples from experiment described in sup.Figure1. Gene expression was compared for the four treatment conditions ("virus UV" in red, "free virus UV" in black, FSW in blue and poly(I:C) in green). Mann-Whitney T test, **pvalue < 0,01; *** pvalue<0,005, ****pvalue<0,0001, ns- non significative (n=10).*

#### 2.2 Supplementary Figure 2

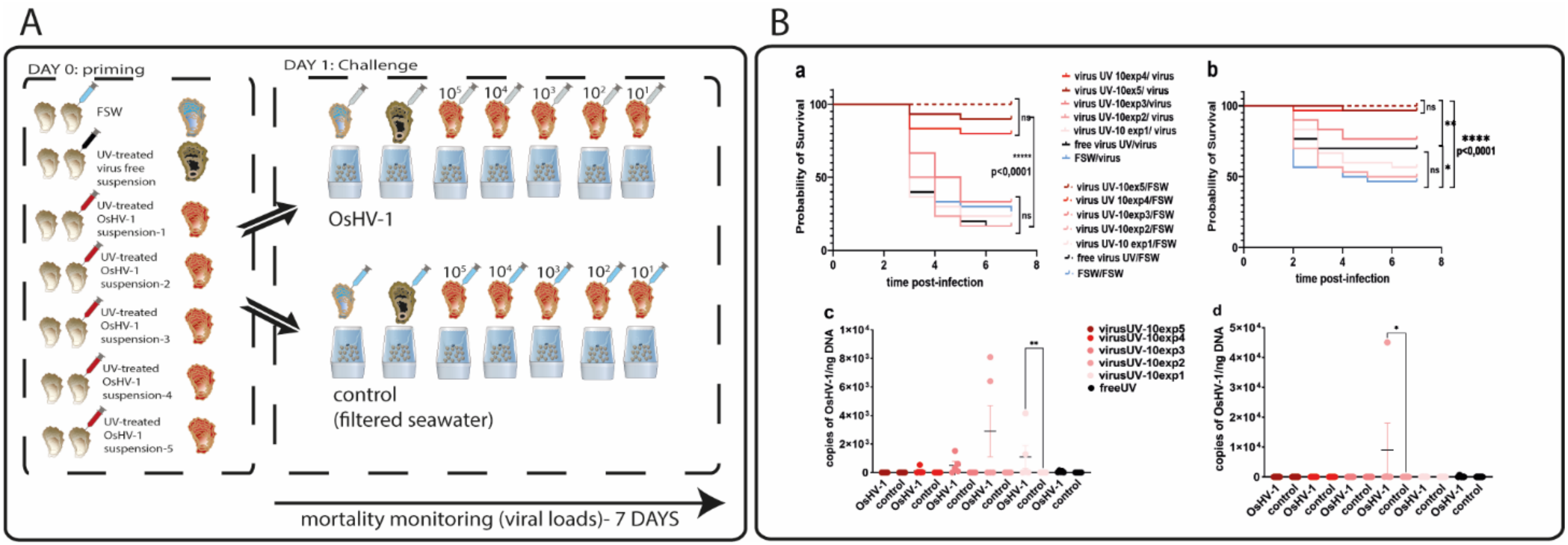


***Sup.Figure 2. Optimal dose of UV-inactivated OsHV-1 suspension treatment***

A. ***Experimental design testing optimal doses of UV-inactivated OsHV-1 suspension treatment.*** *In two independent experiments two different families of OsHV-1 susceptible oysters, F15(a-4 month old) and 01/2020 (b-11 month old), were first injected with 5 different UV-treated OsHV-1 suspension serially diluted after UV-treatment and ranging from $10^5$ UG/µL to $10^1$ UG/µL (10 fold dilutions) or UV-treated virus free suspension and filtered seawater as a negative controls (n=60 for each treatment). 24 hours after the first injection, each treatment group was separated in two groups that were challenged by injection of 100 µL (3,57.$10^4$ copies UG/µL) of OsHV-1 suspension ("virus" condition, n=30) or injected with 100µL of filtered artificial seawater as a control ("SW", n=30). OsHV-1 suspension was prepared as previously described (26). Mortalities were monitored daily for 7 days after challenge. Oysters were sampled for viral DNA, at the end of the experiment.*

B. ***Survival monitoring for optimal dose of UV-inactivated OsHV-1 suspension treatment. a, b. Kaplan-Meier curves*** *showing probability of oyster survival in two independent experiments (a and b) after a primary exposure to different doses of UV treated OsHV-1 (virus-UV in different shades of red), UV-treated virus free suspension (free virus UV, in black) or FSW (blue) and a secondary exposure to OsHV-1 suspension (plain lines) or filtered seawater (FSW, dotted lines). Controls reaching 100% survival* *(oysters challenged with FSW) appear hidden and merged behind the non-treated control line. Results show that injection of UV-treated OsHV-1 increased significantly oyster survival (p<0,001) after challenge for the two most concentrated doses ($10^5$ and $10^4$ copies of OsHV-1/µL) with survival rates of 90% and 80%, respectively (a). This result was confirmed in a similar experiment using a different oyster family where survival rates reached 97% for the two most elevated doses (b). Mortalities in each group of 30 oysters (15 per tank) were monitored for 7 days after infection. **** indicates p-value <0.0001; log-rank test (n=30).* ***c,d. Viral DNA loads in surviving oyster after optimal dose testing.*** *DNA loads monitoring in two independent experiments (c and d ) after challenge with live OsHV-1 were estimated by real time PCR in 10 individual oysters for each condition (different doses of UV-inactivated virus in different shades of red, UV-treated free virus suspension in black) 7 days after challenge with OsHV-1 suspension (OsHV-1) or filtered seawater (control). DNA loads are expressed as copies of OsHV-1/ ng of oyster DNA. Analysis of DNA loads in survival oysters revealed low or undetectable virus DNA.Mann-Whitney T test, * pvalue < 0,05, **pvalue<0,01, ns- non significative (n=5).*

#### 2.3 Supplementary Figure 3

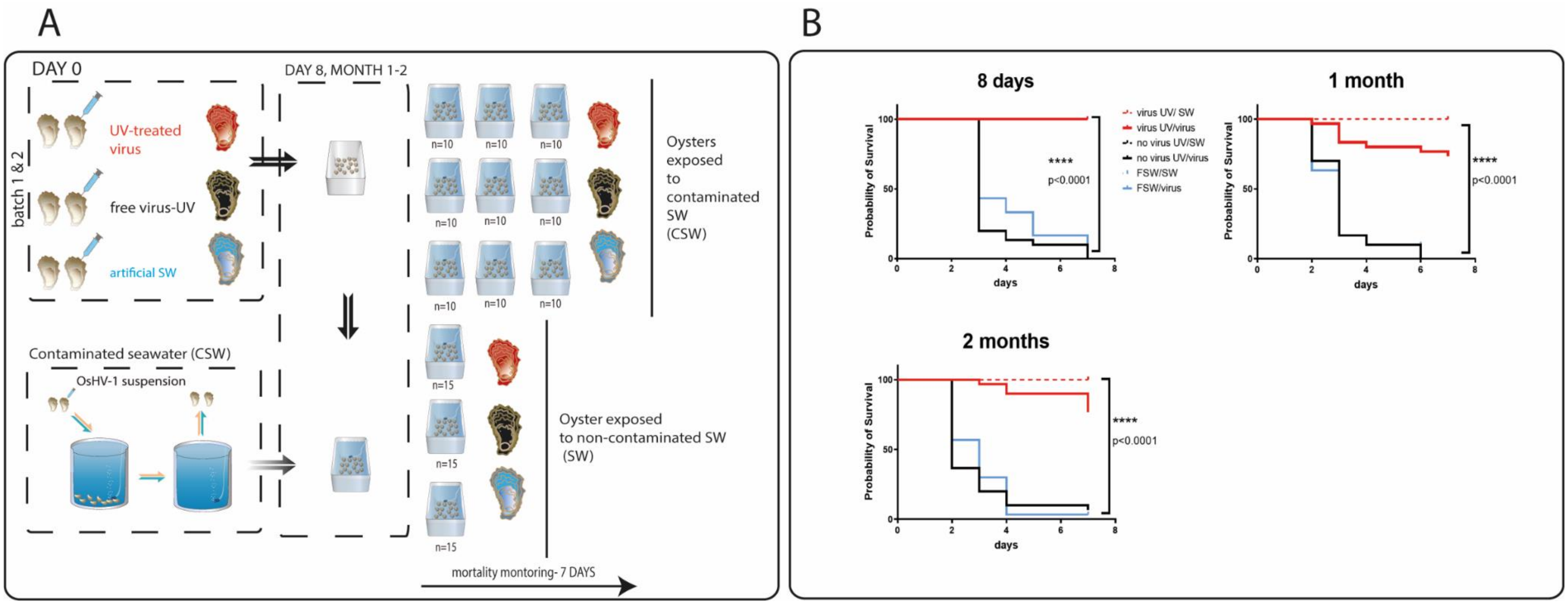


***Sup.Figure 3. UV-inactivated OsHV-1 treatment induces a long-term protection against OsHV-1 infection.***

A. ***Experimental design testing optimal dose of UV-inactivated OsHV-1 suspension treatment.*** *OsHV-1 susceptible oysters (SC18, 8 month old) were first exposed by injection to UV-treated OsHV-1 (virus-UV in red), UV-treated virus free suspension (in black) or FSW (blue) 8 days, 1 month, 2 months or 3 months before being challenged with contaminated seawater (CSW) or filtered seawater as a control (SW). Mortalities were monitored daily for 7 days after the challenge.*

B. ***Kaplan-Meier curves showing probability of oyster survival in an independent experiment (replicate of experiment in Figure 4).*** *After a primary exposure to UV-treated OsHV-1 (virus-UV in red), UV-treated non-viral suspension (free virus UV, in black) or FSW (blue) and a secondary exposure to OsHV-1 contaminated seawater (CSW) (plain lines) or filtered seawater (FSW, dotted lines). Controls reaching 100% survival (oysters challenged with FSW) appear hidden and merged behind the virus-UV/virus or virus/SW condition lines. Mortalities in each group of 30 oysters (15 per tank) were monitored daily for 7 days after infection. **** indicates p-value <0.0001; log-rank test (n=30).*

#### 2.4 Supplementary Figure 4

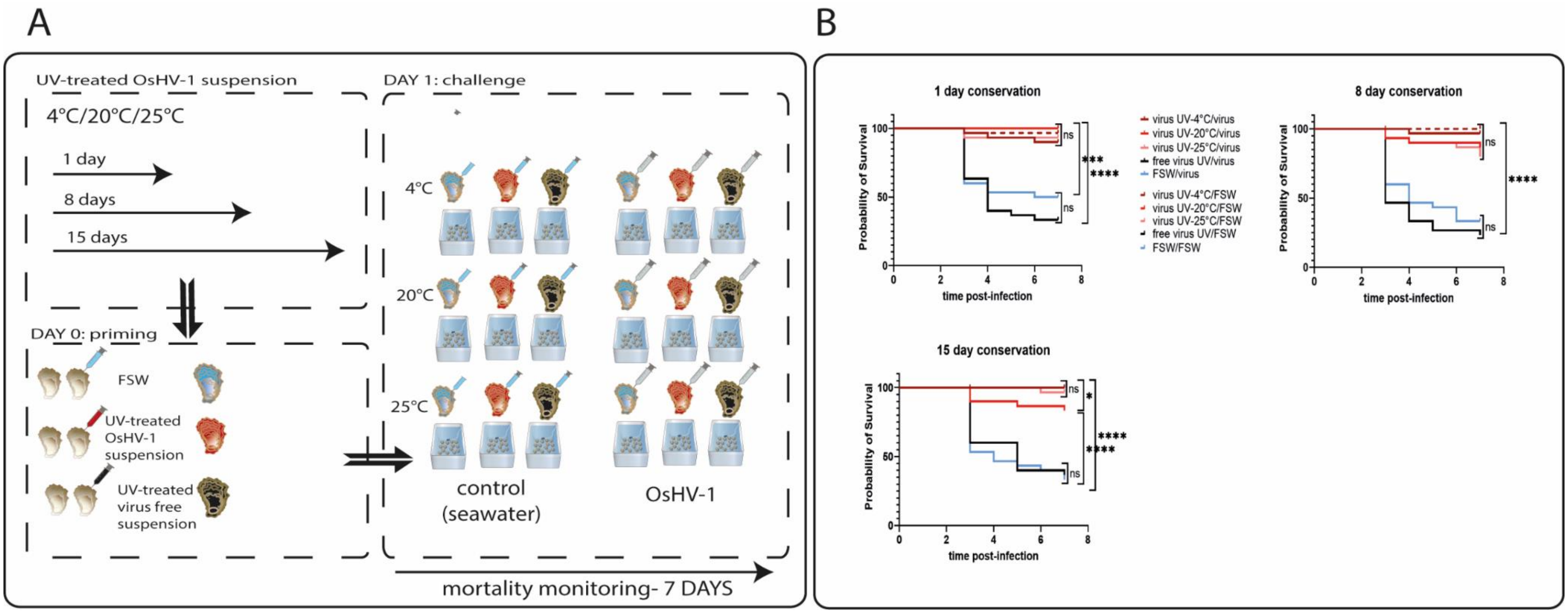


***Sup.Figure 4. Stability of UV-inactivated OsHV-1 suspension.***

***A. Experimental design testing the stability of UV-inactivated OsHV-1 suspension.*** *To test the stability of the UV-treated OsHV-1 suspension overtime, we used a susceptible oyster family (F15, 5 month old). The UV-treated OsHV-1 suspension was conserved for 1 day, 8 days or 15 days at 3 different temperatures (4°C, 20°C, 25°C) before injection into oysters (n=60/condition). UV-treated virus free suspension and filtered seawater were used as negative controls. Oysters were challenged by injection of 100 µL (3,57.10$^{4}$ copies UG/µL) of OsHV-1 suspension ("virus" condition, n=30) or injected with 100µL of filtered artificial seawater as a control ("SW", n=30). Mortalities were monitored daily for 7 days after the challenge.*

***B. UV-treated OsHV-1 suspension is stable aver time and at different temperatures.*** *Kaplan-Meier curves showing probability of oyster survival after a primary exposure to UV treated OsHV-1 (virus-UV in different shades of red) conserved 1 day, 8 days or 15 days at 3 different temperatures (4°C, 20°C, 25°C), UV-treated non-viral suspension (free virus UV, in black) or FSW (blue) and a secondary exposure to OsHV-1 suspension (plain lines) or filtered seawater (FSW, dotted lines). Controls reaching 100% survival (oysters challenged* with FSW) appear hidden and merged behind the “virus UV” condition line. Survival rates indicate that UV-treated OsHV-1 suspensions were highly stable at the three tested temperature and for at least 15 days, efficiently protecting the oysters against OsHV-1. Survival rates were significantly higher in the UV-treated OsHV-1 injected animals at all times, with a mean of 94% (1 day, 15 days) and 89% (8 days). Mortalities in each group of 30 oysters (15 per tank) were monitored daily for 7 days after infection.*pvalue<0,05, ***pvalue<0,005 ,**** pvalue <0.0001 using log-rank test (n=30).
